# A fully 3D-printed optical microscope for low-cost histological imaging

**DOI:** 10.1101/2024.12.16.628684

**Authors:** Jay Christopher, Rebecca Craig, Rebecca E. McHugh, Andrew J. Roe, Ralf Bauer, Brian Patton, Gail McConnell, Liam M. Rooney

## Abstract

We present the manufacture and characterisation of a fully 3D printed, low-cost optical microscope using both a 3D printed chassis and 3D printed illumination and imaging optics. The required commercial components, consisting of a basic camera for image acquisition and light emitting diode controlled by a Raspberry Pi for illumination, are integrated into the 3D printed microscope with the full design shown for ease of replication. Our 3D printed microscope uses a single 3D printed objective lens with a 2.9x magnification and a numerical aperture of 0.07. To benchmark the imaging performance of the system, we used standard test targets and histological specimens, namely, a Giemsa-stained blood smear sample and a thin section of mouse kidney stained with Haemotoxylin and Eosin. We demonstrated that sub-cellular resolution was obtained, and we corroborated this by imaging individual red blood cells and intricate anatomical details of the stained mouse kidney section. All of this was achieved using entirely 3D printed hardware and optics, at a fraction of the cost of a commercial brightfield microscope, while presenting remarkable potential for customisation and increased accessibility for diagnostic imaging applications.

## Introduction

Optical microscopy has long been a cornerstone of scientific discovery, driving advances across diverse disciplines from biology and healthcare to materials science, petrochemistry and geology. Traditional optical microscope instrumentation, while powerful, often comes with substantial costs and infrastructural requirements, limiting accessibility in resource-constrained settings. Recent advances in 3D printing, combined with integration of low-cost, high-performance single board computers (e.g., Raspberry Pi) and low-cost CMOS cameras, have increased democratisation of scientific tools, enabling the development of low-cost microscopes that can be easily customised and distributed^1–6^.

The OpenFlexure Microscope (OFM) represents a significant step in this democratisation, leveraging 3D printing to construct a modular, open-source microscope platform^6,7^. The OFM achieves high spatial resolution and precise sample positioning through its flexure-based mechanical design, enabling high-accuracy imaging at a fraction of the cost of conventional microscopes^7^. Its modular architecture, built entirely from 3D-printed components, supports seamless integration of additional functionalities such as motorised stages and illumination modules for different imaging modalities^8,9^. However, existing implementations of the OFM rely on conventional glass objective lenses, which can cost up to several thousand pounds and be easily damaged. This dependence on commercial optical components limits the accessibility and adaptability that a fully 3D printed microscope could achieve.

Recently, 3D printing technologies have enabled the fabrication of optical-quality lenses suitable for microscopy applications^10,11^. These low-cost lenses have achieved sub-cellular resolution in both brightfield transmission and widefield epifluorescence imaging when used as imaging optics^12^, with images of specimens such as onion cells, cyanobacteria, and plant tissue demonstrating their potential as affordable alternatives to traditional glass optics.

Here we report the first fully 3D printed microscope, bringing together printed components for both the optical and mechanical parts of the system. By combining proven optical designs with additive manufacturing, this study sets a precedent for the development of self-sufficient scientific tools and the potential for rapid, field-customised optical microscopes. We present the design and characterisation of the microscope optics and chassis while demonstrating the potential of the system for accessible low-cost healthcare diagnostics by applying the system to different histological specimens. By demonstrating the viability of an entirely 3D printed microscope, this work further emphasises the transformative potential of additive manufacturing in the field of optical microscopy.

## Methods

### Production of a 3D Printed OpenFlexure Chassis

The OpenFlexure project offers many variations of microscope chassis design which can be customised to the application or needs of the user. For this study, we constructed the 3D printed microscope chassis from the OpenFlexure v6 design, which was suited to the inclusion of 3D printed optics, as opposed to conventional RMS threaded optics and camera adaptors used by other OpenFlexure models. Moreover, we did not require the more recent OpenFlexure design variations which include advanced features such as motorised stage scanning or a specimen riser. .STL files were obtained from the OpenFlexure project website^13^ and processed in Bambu Studio (v1.10.1.50; Bambu Lab, China), where the printing parameters were set for PLA filament (Basic Black; Bambu Lab, China). The layer height was set to 0.2 mm, with an infill density of 15%, and the chassis was printed with no requirement for additional supports. A fused deposition modelling printer (X1C; Bambu Lab, China) was used with a 0.2 mm nozzle, which resulted in a total print time of 10 hours, using 260.9 g of filament (total manufacturing cost = £6.52). The printed microscope chassis was assembled as per the OpenFlexure assembly guide for the ‘Basic Optics’ model, except for the fitted optics and detector which we describe below.

### Production of a 3D Printed Objective Lens and Condenser Lens

The microscope condenser lens design was acquired from a commercial lens manufacturer. We chose the prescription of a 12.7 mm diameter plano-convex lens with a focal length of 35 mm (37791, Edmund Optics, USA); no changes to the geometry of the lens form was required as the refractive index of the print polymer was previously measured to the be the same as BK7 glass^11,14^, which many commercial lenses are made of. The .STEP lens file was imported into Fusion 360 (v2.0.16985; Autodesk, USA) and the polygon count was increased to the maximum available before exporting the model as a .STL file. 3D print files were then generated by importing the .STL file in LycheeSlicer (v5.2.201; Mango3D, France), encoding the print parameters, and exporting as a .CTB file. The condenser lens was printed using a Mars 3 Pro 3D printer with photopolymerising clear resin (RS-F2-GPCL-04; Formlabs, USA) using an exposure setting of 9 seconds for a 10 µm layer height. The printed condenser lens was post-processed by washing with neat isopropyl alcohol for 9 minutes, drying using an air duster (CA6-EU; Thorlabs, USA), and curing for 15 minutes using 385/405 nm light in a curing station (MercuryX; ELEGOO, China). The lens surface was rendered smooth for imaging by spin coating a thin layer of liquid clear resin (version 4; VidaRosa, China) for 10 seconds at 2000 rpm, and post-curing for 10 minutes. The planar surface of the condenser lens was processed by spin coating a thin layer of clear resin (RS-F2-GPCL-04; Formlabs, USA) onto a clean microscope slide for 10 seconds at 2000 rpm and placing the planar surface in contact with the thin resin film. The lens-slide combination was then placed in a vacuum chamber (2L; Bacoeng, USA) fitted to a vacuum pump (Capex 8C; Charles Austen Pumps Ltd, UK) and held under a vacuum of 90 kPa for 30 minutes before curing for 20 minutes as above. The melded lens assembly was then frozen at -20°C for 3 minutes, and the lens was carefully levered from the slide to produce the final lens element, as described previously^11,12^.

The microscope objective lens design was acquired from a commercial manufacturer and, as described above, converted to a .CTB 3D print file. A plano-convex lens with 20 mm focal length and 12.7 mm diameter (LA1074; Thorlabs, USA) was selected, providing a non-infinity design with a 2.9× theoretical magnification when placed at 20 mm from the specimen plane. The objective lens was printed using a Mars 2 3D printer (ELEGOO, China) with 10 µm layer height and an exposure time of 9 seconds using a photopolymerising clear resin (RS-F2-GPCL-04; Formlabs, USA). The objective was then washed using 100% isopropyl alcohol for 9 minutes and dried using an air duster (CA6-EU; Thorlabs, USA), before being post-processed as described for the condenser lens above to produce imaging quality surfaces^11,12^. The objective lens design was modified to include a 25.4 mm diameter, 2 mm thick ring around the lens, which provided the means to fit the lens to the microscope chassis for imaging.

### Specimen Preparation

A blood smear specimen was prepared by collecting blood from a healthy volunteer and smearing along the length of a clean microscope slide. The blood film was air dried before immersing in a bath of 100% methanol, further air drying, and covering with a 5% (w/v) solution of Giemsa stain. The specimens were left to stain for 30 minutes before being rinsed thoroughly with tap water and air dried prior to imaging.

Mouse kidneys were harvested and washed three times in phosphate buffered saline before being fixed in 5 ml of 10% (v/v) neutral buffered formalin (HT501128; Merck, USA) overnight at 4°C. Fixed samples were transferred to 70% (v/v) ethanol and thin sectioning was performed by the University of Glasgow Veterinary Pathology Laboratory. Kidney specimens were processed in ethanol and xylene before embedding in paraffin wax and 2 µm-thick sections were cut and incubated on slides at 60°C prior to staining. Sections were then rehydrated through an alcohol gradient and incubated in haematoxylin (GHS132; Merck, USA) for 5 minutes. Slides were washed with water and differentiated in 1% acid alcohol before washing again. Putts Eosin staining (RBB-0100-00A; CellPath, UK) was conducted for 5 minutes prior to a final wash step. The stained sections were dehydrated through an alcohol gradient and mounted in Histo-Clear (HS2001GLL; National Diagnostics, USA).

### Brightfield Transmission Imaging with a Fully 3D Printed Microscope

The fully 3D printed microscope was assembled by push fitting the 3D printed condenser lens into the OpenFlexure condenser assembly, and placing the 3D printed objective lens into the OpenFlexure objective holder before securing it by screwing a retaining ring around the lens raft. A colour CMOS camera (CS165CU(/M); Thorlabs, USA) was used as a detector, and this was placed at 50 mm from the objective lens, in the base of the OpenFlexure chassis. A small white light emitting diode (LED) (NSPW515DS; RS Components, UK) was used as an illumination source, and this was secured at the top of the OpenFlexure condenser assembly at a distance of approximately 30 mm from the 3D printed condenser lens. The LED light source was triggered using a Raspberry Pi 4 (Raspberry Pi, UK). Camera acquisition and image processing was performed using an Elitebook 840 G7 (Hewlett–Packard, USA) running a 64-bit Windows 10 Enterprise operating system with an Intel Core i5-10310U 1.70 GHz quad-core processor with 16 GB of 2666 MT/s DDR4 RAM. Image acquisition was completed using ThorCam (64-bit, v3.7.0; Thorlabs, USA). An exploded view of the fully 3D printed microscope is presented in Figure 1.

**Figure 1.**
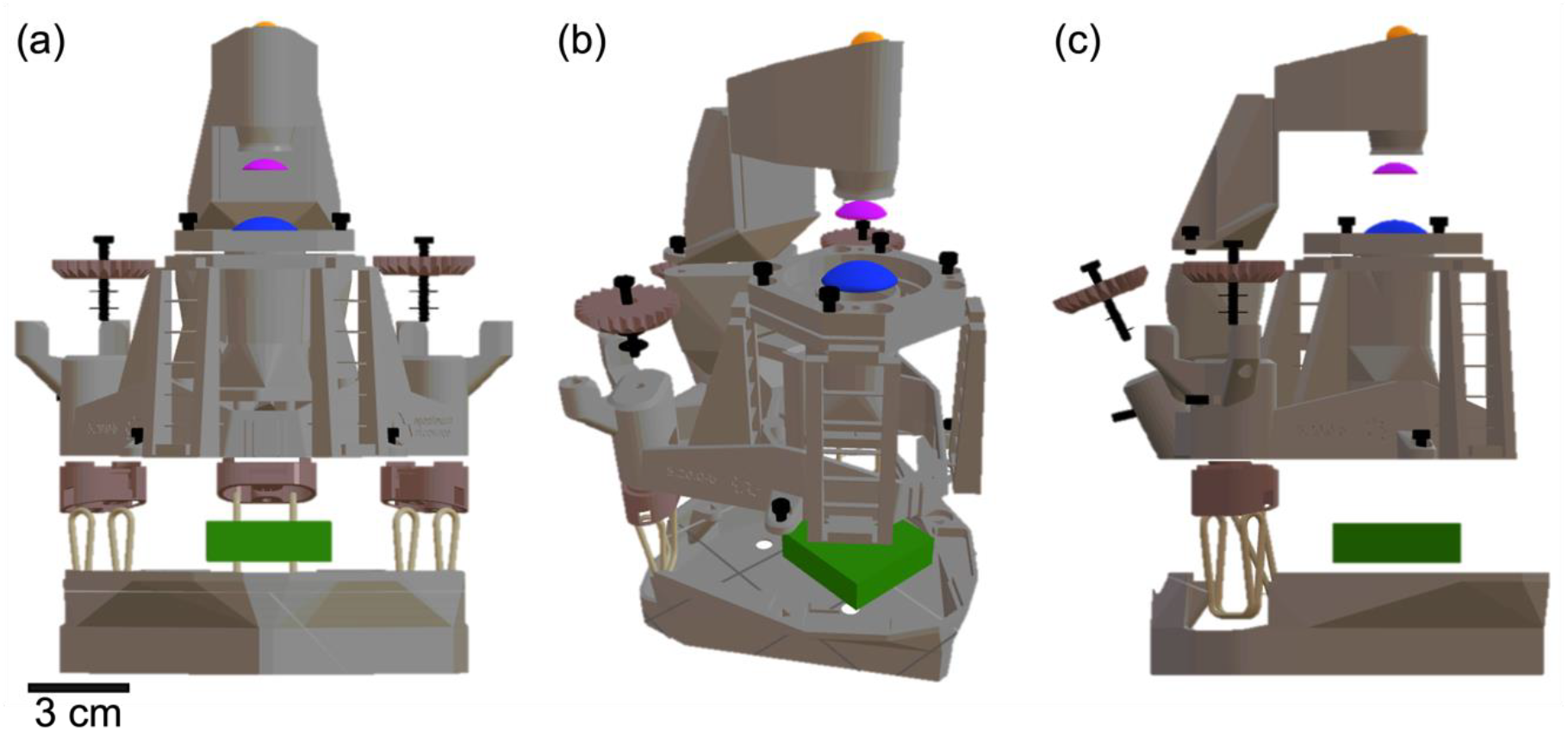
An exploded view of a fully 3D printed microscope. A front-facing **(a)**, oblique **(b)**, and side-facing view **(c)** of the microscope are presented, with the chassis coloured in grey and the positions of additional components highlighted with different colours (orange = LED light source, magenta = 3D printed condenser lens, blue = 3D printed objective lens, green = CMOS camera). A scale bar is presented in (a) to provide a visualisation of the size of the microscope setup.

### Calculating the Magnification and Numerical Aperture of a Fully 3D Printed Microscope

We calculated the Numerical Aperture (*NA*) of the imaging system by applying the Rayleigh criterion in the context of our experimentally determined resolution limit, *dx*,*y*, where the wavelength (*λ*) was assumed as 550 nm (Equation 1)^15^.

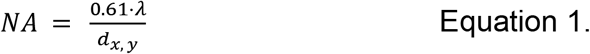

We calculated the magnification (*m*) based on the relationship between the diameter of the field of view (*FOV*) and the width of the camera sensor (*H*), given by Equation 2.

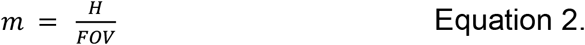

To measure the spatial resolution of the imaging system, we imaged a United States Air Force (USAF) resolution test target (2015a USAF; Ready Optics, USA) using the brightfield transmission setup described above. To measure the size of the field of view, we imaged a 10 mm stage micrometer graticule specimen (R1L3S1P; Thorlabs Inc., USA) using the same setup.

### Image Processing and Analyses

All image processing and analyses was conducted using FIJI (v1.54f)^16^. Spatial resolution and field of view dimensions were measured from the USAF test target and graticule images, respectively. A line intensity plot profile was acquired for the measurement region of each specimen as described in the Results section and used to determine the image scaling parameters.

Chromatic aberrations in the raw image data were corrected using the *Correct 3D Drift* plugin in FIJI. Briefly, the raw RGB image was split into three separate colour channels and recompiled as image stack, where each image was reassigned from a colour to a time point. Correction was limited to only *x, y* drift and the performance of the correction was verified by comparing line intensity profiles of the same region of interest (ROI) in the raw and corrected datasets.

## Results

### 3D printed optics provide a fully 3D printed microscope with cellular resolution

The spatial resolution of the fully 3D printed microscope was determined by imaging a USAF resolution test target with line spacing ranging from 31 µm to 137 nm. Figure 2a shows a raw image of the test target, with a magnified ROI of the smallest resolvable structures, corresponding to Group 6. The image shows clear chromatic aberrations, which were digitally corrected post-acquisition (shown in Figure 2b). A line intensity plot profile was measured across Group 6, Elements 2-6, for both the raw and chromatically corrected data and presented in Figure 2c. The post-acquisition correction provided a marginal increase in contrast without altering the relative position of the image features, with a resolution limit of Group 6, Element 5 corresponding to 4.922 µm as defined by the manufacturer. By applying the Rayleigh criterion, as per Equation 1, we calculated the NA of the system to equal 0.07.

**Figure 2.**
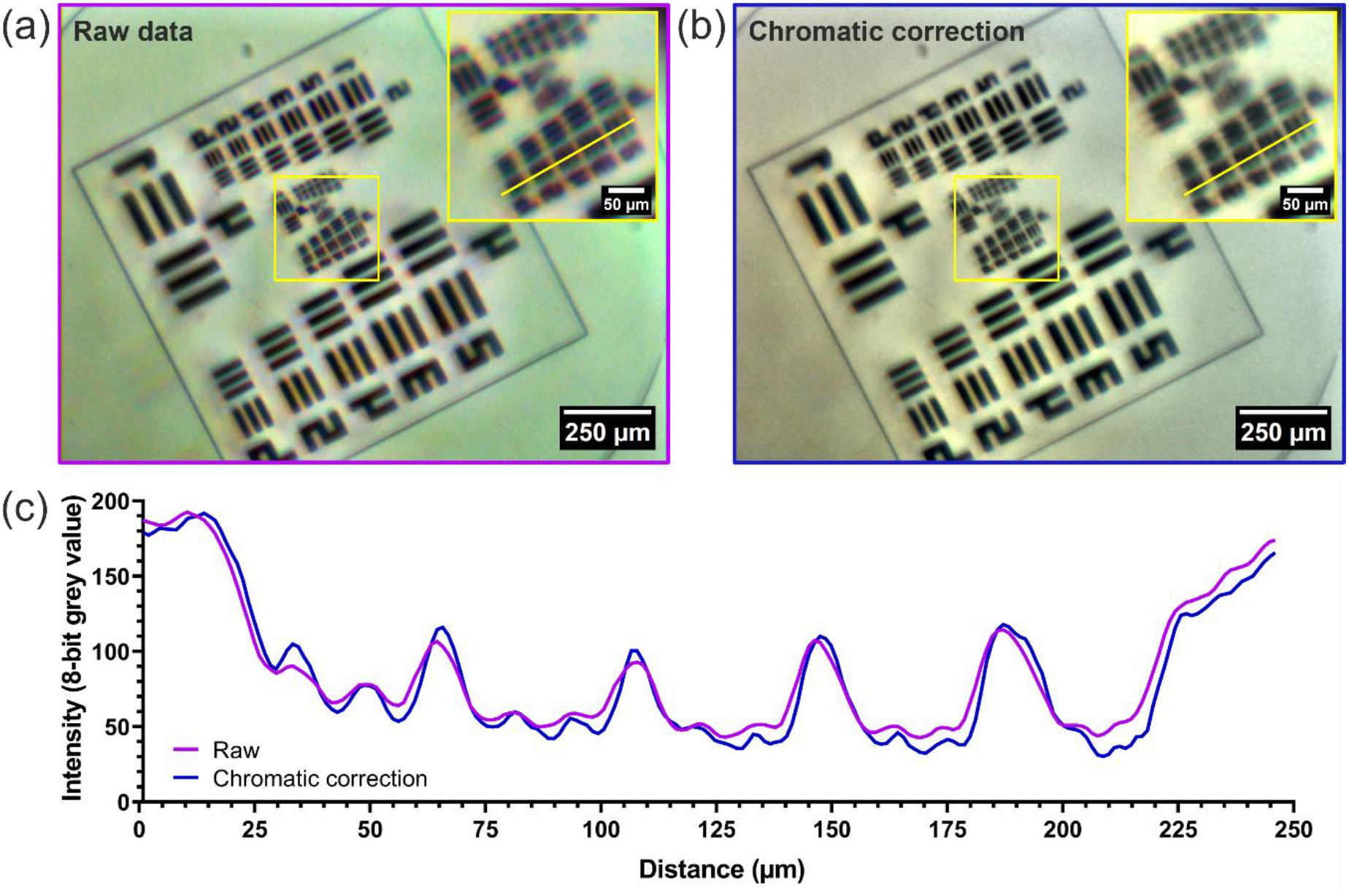
Measuring the resolution of a fully 3D printed microscope post-acquisition chromatic correction. An image of a USAF resolution target imaged using a fully 3D printed microscope before **(a)** and after digital chromatic aberration correction **(b)**. A magnified region of Group 6 is shown for each image, where a linear intensity plot profile was measured through Group 6, Elements 2-6. **(c)** An intensity plot profile of the raw and chromatic correction data is presented. The simple process of correcting chromatic aberrations provided a resolution of 4.9 µm (Group 6, Element 5) with higher contrast than the raw data.

### Determining the magnification and assessing illumination homogeneity of a fully 3D printed microscope

A 10 mm graticule was used to determine the 3D printed microscope scaling parameters and to provide a visualisation of the contrast across the field of view. Figure 3a shows a chromatically corrected image of the graticule, with a line intensity plot profile presented in Figure 3b. We observed reasonable contrast across the field of view and homogeneity of illumination with a decrease of 24% towards the edge of the measurement region (see Figure 3b), permitting the detection of graticule increments with a 50 µm spacing. From this, we measured the attainable field of view for our fully 3D printed microscope to be approximately 1.7 mm. Using Equation 2, we calculated the effective magnification of the imaging setup to equal 2.90×.

**Figure 3.**
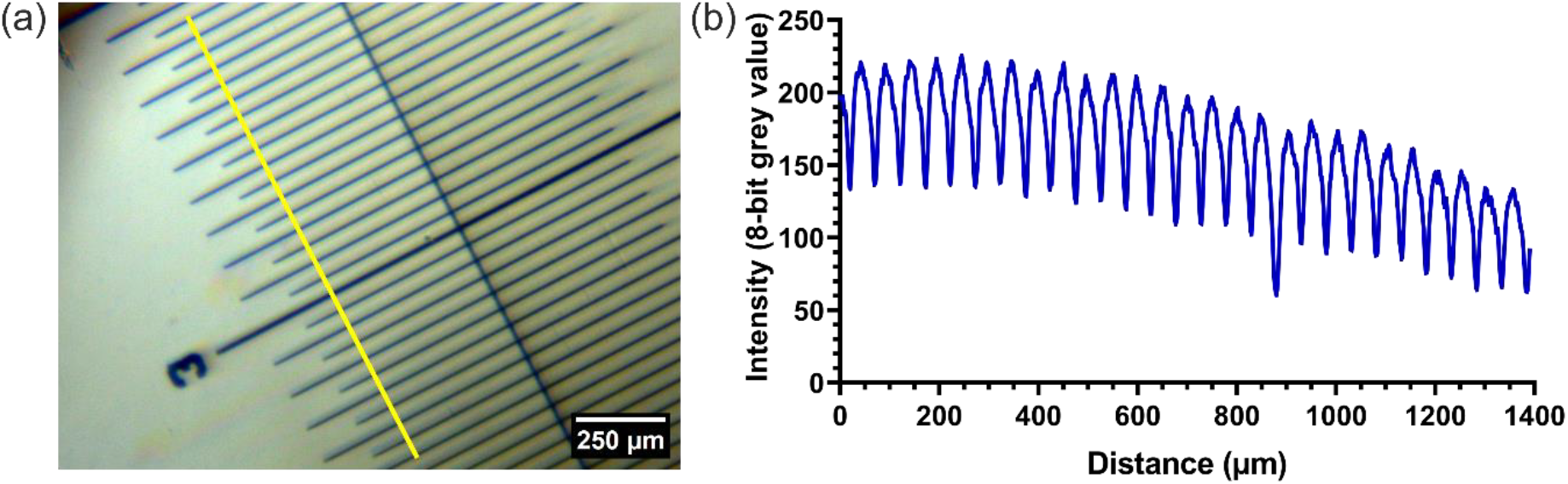
Demonstrating the homogeneity of illumination and determining the magnification and scaling parameters for a fully 3D printed microscope. **(a)** A brightfield transmission image of a 10 mm graticule obtained with a fully 3D printed microscope, where major increments are separated by 100 µm. **(b)** An intensity plot profile measured from the linear ROI presented in (a). The plot shows good contrast over the measurement region, with clear visualisation of the graticule structure.

### Exploring the potential of a fully 3D printed microscopy for histological imaging

We demonstrated the potential of a fully 3D printed microscope for histological imaging using two routine histopathology specimens, namely a blood smear and a thin tissue section (Figure 4). Figure 4a shows an image of a Giemsa-stained blood smear with two magnified ROIs from different regions of the FOV. Both regions exhibit high contrast and clearly resolved individual blood cells using a simple objective lens comprised of a single 3D printed lens. Moreover, the two regions show a similar focus, suggesting that despite the sphericity of the imaging lens that a sufficiently flat field is produced to image at this low magnification. Figure 4b shows an image of an H&E-stained thin section of mouse kidney. Large anatomical structures such as in interlobular arteriole containing red blood cells can be clearly observed in the centre of the FOV (shown by the white arrow in Figure 4b), with additional structures visible in the magnified ROI. Here, the fully 3D printed microscope facilitates visualisation of intricate anatomical details, such as arrangements of nephrons in a medullary ray spanning the corticomedullar junction of the kidney.

**Figure 4.**
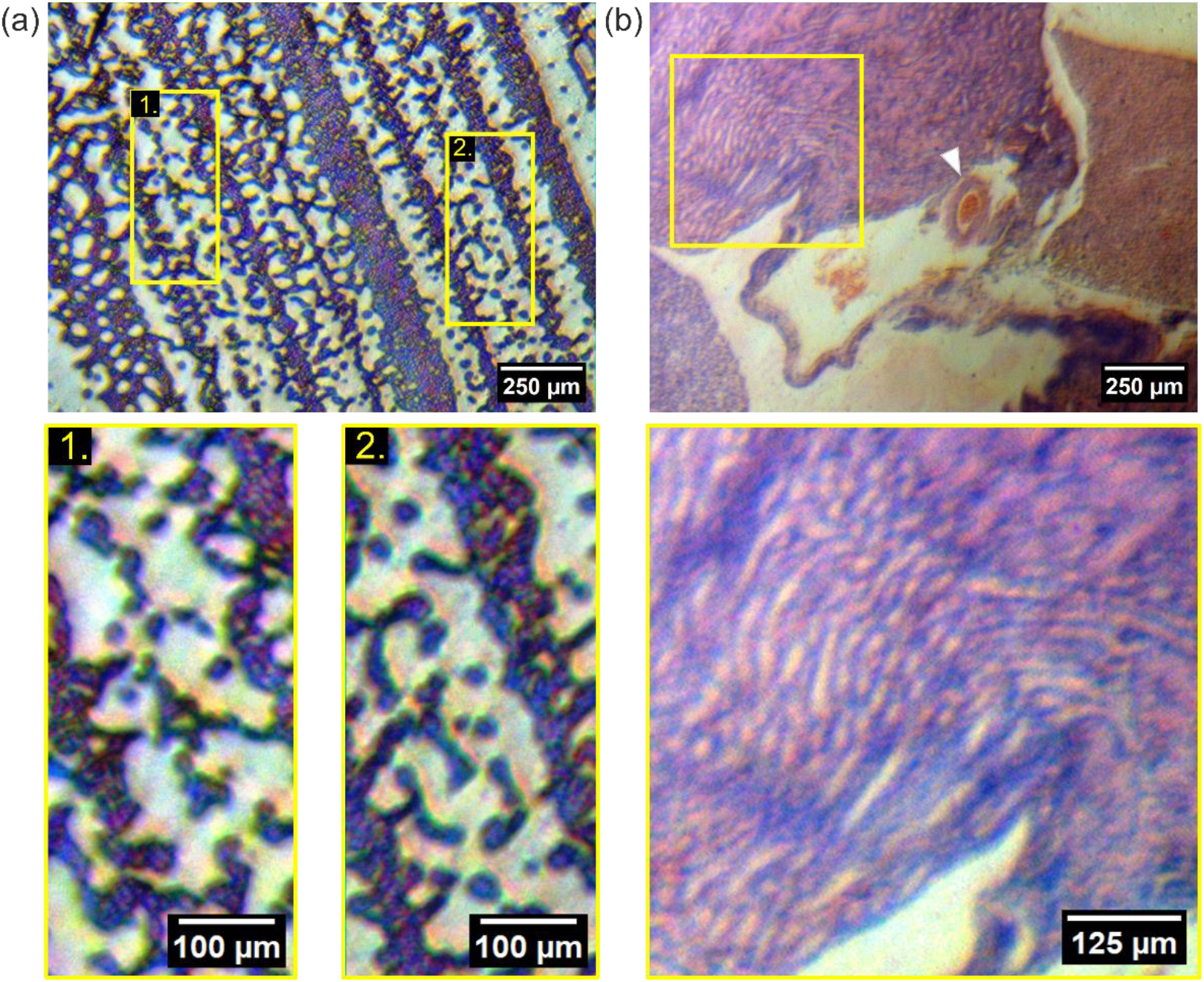
Implementation of a fully 3D printed microscope for low-cost histopathology imaging. **(a)** An image acquired of a Giemsa-stained blood smear. Two ROIs are shown, where individual red blood cells can be resolved over the 1.7 mm field of view. **(b)** An image of a H&E-stained mouse kidney. The thin section shows structures such as an interlobular arteriole (white arrow) and renal tubules, with a magnified ROI showing the organisation of nephrons in a medullary ray spanning the corticomedullary junction.

## Discussion

We present the first fully 3D printed microscope where both the microscope chassis and the illumination and imaging optics are entirely 3D printed. We combined the well-established and proven design of the OpenFlexure microscope body with our previously characterised 3D printed lenses, creating a fully customisable, fully open source, low-cost imaging system capable of resolving individual mammalian cells. We demonstrated the use of the fully 3D printed system to routine histology specimens, a blood smear and a thin stained tissue section, providing encouraging results for diagnostic imaging applications in low-resource settings using an entirely home-manufactured microscope.

We first characterised the performance of our 3D printed microscope by measuring the resolution and magnification of the system. The 3D printed lens prescriptions we used provided a total magnification of 2.9×, producing an imaging field of view equal to 1.7 mm, with single-cell spatial resolution on the order of 5 µm. The performance of our 3D printed microscope is comparable to low-cost systems such as the Foldscope and Enderscope. While the Foldscope, a paper origami-style microscope, can obtain sub-micron resolutions in comparison to our fully 3D printed microscope^17^, our filament-printed chassis in conjunction with lab-manufactured optics provides significant potential for rapid customisation across multiple modalities at low cost while maintaining a high standard of axial and lateral translation accuracy. The use of a ball lens in the Foldoscope provides a magnification range of 50-340×, but introduces edge artefacts and spherical aberrations which result in field curvature and distortion^17^. In contrast, the flexibility of our fully 3D printed microscope system provides the user complete control over objective integration as per the needs of the specimen due to the unique potential for 3D printed objectives to be designed with custom prescriptions and combinations in mind. Compared to the Endoscope, a microscope built from the chassis of a Fused Deposition Modelling (filament-based) 3D printer, the spatial footprint of our fully 3D printed microscope is minimal (15 cm^2^ versus 40 cm^2^, respectively). However, with the addition of post-processing stitching and tiling, the Enderscope can capture significantly larger imaging areas with similar resolution to what we have shown in this study^18^. A key highlight of our fully 3D printed microscope in conjunction with our custom optics is the ease of reproducibility and on-the-field sample study potential due to the portability of our system. An integral comparison is the native OpenFlexure system using commercial optics, which exhibits comparable resolution and image quality to our fully 3D printed system, shown recently when applied to histology imaging of oesophageal biopsies^4^. Moreover, Rosen *et al*. also documented the inherent chromatic aberrations using their OpenFlexure setup as we observed and corrected during our imaging experiments^4^. It is important to note that one could apply this simple ImageJ correction operation to any RGB image to mitigate these intrinsic aberrations.

Although we have demonstrated a fully 3D-printed microscope using the OpenFlexure frame, a fully 3D-printed microscope could leverage alternatives to OpenFlexure, such as the UC2 (You See Too), MicroscoPy or the MultiModal Modular Microscopy for All (M4All) platforms, to enhance modularity and accessibility. UC2’s versatile LEGO-inspired design allows users to build and customise microscopy setups, including fluorescence and polarisation configurations, using easily printable components^2^. The MicroscoPy platform, built from a combination of LEGO bricks, 3D printed parts and commercial optics, provides another route for flexibility by relying on the myriad potential forms that LEGO builds can afford^19^. Moreover, the modularity of the M4All platform, which uses arrays of 3D printed cubes that assemble into different functional components^20^, would expand the number of imaging modalities available from a fully 3D printed microscope. By combining these systems with 3D printed optics, researchers and educators could develop cost-effective, scalable, and robust tools for diverse imaging applications, from histopathology and biological research to education.

We also consider the commercial alternative, where a research or clinical grade microscope can be upward of £15k and, in the case of field diagnostics in low-resource settings, presents issues surrounding the availability of instrument servicing and replacement optics or optomechanical parts. As such, building on our previous work in additive manufacturing and characterising high-performance 3D printed lenses for optical microscopy^11,12^, our contribution of a fully 3D printed microscope provides the benefit of *ad hoc* customisation which directly addresses key issues around component supply that often impede open-source microscopy initiatives and field diagnostics in low-resource settings. Our fully 3D printed, open-source system is achievable using high-precision, low-cost, consumer-grade desktop printers with material costs amounting to £7.00 per microscope before the inclusion of a light source and camera.

For brightfield imaging, the illumination should be homogeneous across the field of view, otherwise aligned for Köhler illumination. Figure 3 demonstrates that, despite our fully 3D printed microscope obtaining single-cell resolution, we suffered from inhomogeneity of illumination over the field that was difficult to correct at the time of acquisition due to the lack of suitable LED fixings on the illumination arm of the microscope. Since the optical performance of the 3D printed optics used in this study has been validated extensively^11,12^ we are confident that the quality of 3D printed lenses is unlikely to be the dominant source of illumination inhomogeneity we observed. The alignment of the illumination relative to the Raspberry Pi camera in the native OpenFlexure was ancillary to the direct coupling of the camera to the objective housing. However, due to the design of our system where the camera was uncoupled, there was little margin when manually aligning the illumination source to the optical axis. Off-axis optical alignment exacerbated the effects of the chromatic aberrations we observed, but these were easily corrected with negligible impact on the spatial positioning of the image features. A potential solution to rectify the alignment issues would be to modify the printed microscope chassis design to include adjustable field and condenser diaphragms to facilitate Köhler illumination, providing a uniform intensity over the field of view and minimising intensity artefacts in brightfield transmission imaging. Despite the inhomogeneity, our data concluded that the 3D printed lens designs had a total effective magnification of 2.9×, while resolving single cells, which was within the range expected compared to the prescription of the lens and the distance of the objective lens from the detector. Thanks to the flexibility afforded by printing of optical elements, lenses with alternative magnifications and numerical apertures would be possible, and these could be explored. Finally, the visualisation of biological structures using entirely 3D printed hardware and optics demonstrates great potential for simple diagnostic imaging applications and further refinement by the inclusion of additional 3D printed lens elements.

## Acknowledgements

JC was funded by an Engineering and Physical Sciences Research Council (EPSRC) studentship (EP/T517938/1). RC was supported by an EPSRC studentship. REM and AJR were supported by the Medical Research Council (MRC) (MR/V011499/1). RB was supported by the EPSRC (EP/S032606/1) and the Biotechnology and Biological Sciences Research Council (BBSRC) (BB/Z51486X/1). BP was supported by the Royal Society (CHG/R1/170017). GM was supported by the MRC (MR/K015583/1) and the BBSRC (BB/P02565X/1 and BBT011602) LMR and GM were funded by the Leverhulme Trust.

## Data Availability Statement

The data that support the findings of this study are openly available at the University of Strathclyde KnowledgeBase; https://doi.org/10.15129/b4d95965-f421-4396-acfb-11bf1c68b17c.

